# RNA-binding tunes the conformational plasticity and intradomain stability of TDP-43 tandem RNA recognition motifs

**DOI:** 10.1101/2024.02.25.581941

**Authors:** Busra Ozguney, Priyesh Mohanty, Jeetain Mittal

## Abstract

TAR DNA binding protein 43 (TDP-43) is a nuclear RNA/DNA-binding protein with pivotal roles in RNA-related processes such as splicing, transcription, transport, and stability. The high binding affinity and specificity of TDP-43 towards its cognate RNA sequences (GU-rich) is mediated by highly conserved residues in its tandem RNA recognition motif (RRM) domains (aa:104-263). Importantly, the loss of RNA-binding to the tandem RRMs caused by physiological stressors and chemical modifications promotes cytoplasmic mislocalization and pathological aggregation of TDP-43. Despite the substantial implications of RNA in TDP-43 function and pathology, a comprehensive characterization of the effect of RNA-binding on conformational dynamics, interdomain interactions and intradomain stability of the tandem RRMs has not yet been conducted. Here, we employed all-atom molecular dynamics (MD) simulations to assess the effect of RNA-binding on the conformational landscape and intradomain stability of TDP-43 tandem RRMs. Our simulations reveal a high intrinsic conformational plasticity of the tandem RRMs in the absence of RNA which surprisingly, is accompanied by a tendency of RRM1 to adopt partially-unfolded conformations. While binding to RNA limits the overall conformational space of the tandem RRMs and promotes intradomain stability, several RRM-RNA contacts mediated by highly conserved residues are observed to be far more dynamic than previously inferred from NMR structural ensemble. Overall, our simulations reveal how RNA dynamically tunes the structural and conformational landscape of TDP-43 tandem RRMs, contributing to physiological function and mitigating pathological aggregation.

**SIGNIFICANCE:** The cytoplasmic mislocalization and aggregation of TDP-43 due to loss of its RNA-binding capability is associated with the onset and progression of neurodegenerative diseases such as Amyotrophic lateral sclerosis (ALS) and frontotemporal dementia (FTD). Due to the flexible nature of RNA and the presence of a disordered linker between RRM domains, characterizing the dynamic interactions between RRMs-RNA and/or RRM1-RRM2 by experiments alone has remained challenging. In this study, we performed all-atom simulations initiated from the NMR conformers of RNA-bound tandem RRMs of TDP-43 to investigate their underlying structural and conformational dynamics. Our findings indicate that RNA binding effectively reduces conformational heterogeneity in the tandem RRMs and acts as a protective factor for the unfolding and aggregation of RRM1. These effects are achieved through a combination of stable and dynamic protein-RNA interactions which involve highly conserved amino acids.

## INTRODUCTION

TAR DNA binding protein 43 (TDP-43) is a 414-amino acid multidomain protein characterized by distinct structural elements: an N-terminal domain (NTD, amino acids 1-78) [1], tandem RNA recognition motifs (RRM1, amino acids 106-176) and RRM2, amino acids 193-259) [2, 3], and a prion-like C-terminal domain (CTD, amino acids 264-414). Under normal physiological conditions, TDP-43 primarily localizes to the nucleus [4] and interacts with GU-rich nucleic acid sequences (RNA/DNA) enriched in the target transcripts via the tandem RRMs [5–9]. Ubiquitin-positive, cytoplasmic TDP-43 inclusions were observed in the brains of individuals with amyotrophic lateral sclerosis (ALS) and frontotemporal lobar degeneration (FTLD) [10], suggesting a possible link between TDP-43 mislocalization and neurodegeneration. TDP-43 plays pivotal roles in cellular processes such as gene transcription control, RNA stability maintenance, alternative splicing, RNA translation, and transport [4, 11]. Importantly, the binding of nucleic acids to TDP-43 prevents its mislocalization to the cytoplasm [12], inhibits pathological aggregation [13, 14], and facilitates the formation of functional higher-order TDP-43 assemblies [15].

The N and C terminal domains of TDP-43 contribute towards the formation of the biomolecular condensates through liquid-liquid phase separation (LLPS) [16–24]. Recently, it was demonstrated that RNA-binding maintains the NTD oligomerization crucial for NTD-driven LLPS in the nucleus [25]. Factors such as disease-associated mutations, temperature fluctuations, and exposure to oxidative, osmotic, and chemical stressors [26, 27], can impair nucleic acid binding, disrupt functional LLPS and ultimately cause mislocalization to the cytoplasm. Molecular chaperones known as heat shock proteins (HSPs) have gained increasing attention in regulating the solubility of RNA-free TDP-43 [28–32]. RNA-free TDP-43 interacts with Hsp70 through its RRMs and Hsp90 through its CTD [30]. The interplay between RNA-binding and Hsp70-binding presents a novel perspective on disease progression, as Hsp70 selectively binds to RNA-free TDP-43, stabilizing the Apo state of the protein [28]. The presence of RNA impedes Hsp70 binding, suggesting a role in maintaining the proper fold and potentially preventing misfolding [29]. Therefore, assessing the differences in structural stability of tandem RRMs and the overall conformational landscape in the RNA-free and RNA-bound states becomes important in order to enhance our understanding of TDP-43 function and dysfunction.

Given the critical involvement of RRM domains in mediating pathological TDP-43 aggregation [12, 13, 33-35], a detailed characterization of the tandem RRM conformational ensemble and intradomain stability may provide insights into nature of aggregation-prone conformations and the interactions implicated in the stabilizing effect of GU-rich RNA. However, the presence of a 14 amino-acid long disordered linker connecting the tandem RRMs [2, 36], coupled with the intrinsic flexibility of the unstructured RNA, poses a challenge in characterizing the protein-RNA interface through conventional structure determination approaches such as x-ray crystallography [37, 38]. Alternatively, the integration of knowledge obtained by the nuclear magnetic resonance (NMR) experiments with molecular dynamics (MD) simulations shows promising results in providing a more refined representation of the protein-RNA complexes [37, 39]. While previous studies have utilized MD simulations to investigate the intradomain stability of TDP-43 RRMs and their interactions with RNA [40–44], a comprehensive understanding of the tandem RRM conformational landscape and how RNA modifies it to promote TDP-43 function and inhibit pathological aggregation, is still lacking. To this end, we conducted multi-microsecond long MD simulations of the NMR-derived conformer ensemble of TDP-43 tandem RRMs which was previously determined in the presence of a bound GU-rich RNA fragment [2] to investigate and elaborate on the changes in the interdomain flexibility and intradomain stability which occur upon RNA-binding. Furthermore, we aimed to analyze the dynamic inter-RRM and RRM-RNA interactions at an atomistic resolution to elucidate the intricate details of the protein-RNA interface.

Our simulations indicate a high intrinsic conformational plasticity of the tandem RRMs in the absence of RNA and increased conformational fluctuations at the intradomain-level which is associated with a tendency for RRM1 to adopt partially-unfolded conformations. RNA binding effectively reduces the intrinsic conformational variability of the tandem RRM domains and acts as a protective factor for the unfolding of RRM1 by stabilizing its β4/5 region. Finally, GU-rich RNA displays high sequence specificity toward conserved phenylalanine residues in both RRM domains, while the lysine residues in the RRM1 facilitate the formation of a more dynamic binding interface between RRM1 and RNA. In conclusion, our study offers a comprehensive biophysical characterization of the conformational changes in the TDP-43 tandem RRMs associated with RNA binding, bearing relevance for understanding the relationship between its conformational stability and pathological aggregation [45].

## MATERIALS AND METHODS

### Calculation of Conservation Scores

To evaluate the evolutionary rate of each residue in TDP-43 tandem RRM structure, we employed ConSurf [46]. Sequence of tandem TDP-43 was provided in FASTA format to the webserver, and a ConSurf run was launched with default settings. The algorithm conducted a blast search over UNIREF-90 database to determine the homologous sequences. In this initial search, only 35% similarity was considered. Then, redundant sequences were eliminated using CD-HIT clustering method. Remaining sequences were aligned with the method of Multiple Alignment using Fast Fourier Transform (MAFFT). Phylogenetic tree reconstructed by the generated multiple sequence alignment (MSA). Finally, Rate4Site algorithm calculated the per-residue conservation scores.

### Molecular modeling and simulation protocol

The initial structures for the RNA-bound state were taken from the 20 NMR-solution models of TDP-43 tandem RRMs (PDB: 4BS2) (**Figure S1**) [2]. To model initial structures for the simulations of the RNA-free state, a 12-nucleotide long GU-rich (5’-GUGUGAAUGAAU-3’) RNA was removed. Sequence modifications from GU-rich to (A)_12_ or (U)_12_ were introduced using the “swapna” module of Chimera [47]. Construct ΔRRM1 was generated by deleting residues from G96 to K176, and construct ΔRRM2 was created by deleting residues from K192 to Q269. Additional details can be found in **Supplementary Information Table S1**.

Initial structures (20 for each construct) were prepared for all-atom simulations in Gromacs v2022.3 [48]. Structures were solvated in octahedral water boxes with a box length of 10.5 nm, providing approximately 1.5 nm of box padding. The number density of water molecules was kept constant across all structures of the construct. Na+ and Cl-ions were introduced to mimic salt concentration of 100 mM, along with any additional counter ions, 15 Na+ for ΔRRM1, 8 Na+ for ΔRRM2, 13 Na+ for RNA-bound, and 2 Na+ for RNA-free constructs, necessary to neutralize the systems. The Amber03ws force field was applied for protein topology with the tip4p2005 water model. χOL3 force field was chosen to model RNA topology, and details for this choice could be found in **Supplementary Information** [49–53]. The Gromacs files (top and gro) were converted to Amber inputs (parm7 and rst7) using the “gromber” utility in ParMed [54]. Additionally, hydrogen mass repartitioning to 1.5 amu was performed [55].

Production simulations were performed in Amber22 [55] at 1 bar and 300K following a multi-step equilibration process (**Figure S1**). A minimization run was carried out by utilizing the steepest descent algorithm and the conjugate gradient algorithm, with all non-hydrogen atoms of the protein (and RNA) restrained by a 5 kcal/(mol Å) force constant. The energy-minimized structures underwent heating for 5 ns, with temperature linearly increasing from 0 K to 300 K in the first half and then kept constant at 300 K in the second half. The force constant was reduced to 1 kcal/(mol Å) with a time step of 2 fs in this simulation. Subsequently, two NVT simulations were conducted: a 5 ns equilibration at 300 K with a timestep of 4 fs and another 5 ns equilibrium at 300 K with all restraints released. Systems were further equilibrated for 10 ns with a 4 fs time step under an isothermal-isobaric ensemble (NPT) using a Monte Carlo barostat [56] with an isotropic coupling of 1.0 ps at a pressure of 1 bar. Langevin dynamics with a 1.0 ps^-1^ friction coefficient was employed for temperature control, and bonds involving hydrogen atoms were constrained using SHAKE [57]. A non-bonded cutoff of 0.9 nm was applied for short-range non-bonded interactions, and long-range electrostatic interactions were treated using the Particle Mesh Ewald (PME) method [58]. Finally, systems were subjected to 1 µs NPT production run with the same parameters used for equilibrium.

### Structural and Contact Analysis

All structural analyses, RMSD, dRMSD, RMSF, Rg, were conducted using Gromacs v2022.3 utility programs [48]. Distance-based contact maps were computed using the “neighbor search library” module of the MDAnalysis [59]. A van der Waals contact between two entities (protein residue or RNA nucleotide) was defined if at least one non-hydrogen atom of one entity was within 0.6 nm of a non-hydrogen atom in the other entity.

## RESULTS

Lukavsky et al. [2] solved the solution structure of the tandem RRM domains of TDP-43 spanning amino acids:102-269 in complex with a 12-nucleotide GU-rich RNA molecule (**Figure S1**). RRM1 and RRM2 exhibit a distinct tandem arrangement when compared to other tandem RRM-RNA complexes, e.g., Sex-lethal, PABP, HuD and Nucleolin [2]. Notably, RNA is bound in the reverse direction, rather than in the typical 3’ to 5’ direction to RRM1 and RRM2. TDP-43 RRMs also exhibit a high degree of evolutionary conservation in the residues constituting the RNA binding pocket, as shown by ConSurf scores in **Figure 1A**, which underscores the functional significance of RRM-RNA interactions. The presence of a flexible linker connecting the two RRM domains, however, implies a dynamic conformational ensemble comprising of alternative inter-RRM orientations and interactions. Accordingly, we used multi-replica all-atom molecular dynamics (MD) simulations to characterize conformational landscape of TDP-43 tandem RRMs; both in the presence and absence of GU-rich RNA (**Figure S1**). The aggregate duration of the simulation replicas are 20 µs for each state.

**Figure 1.**
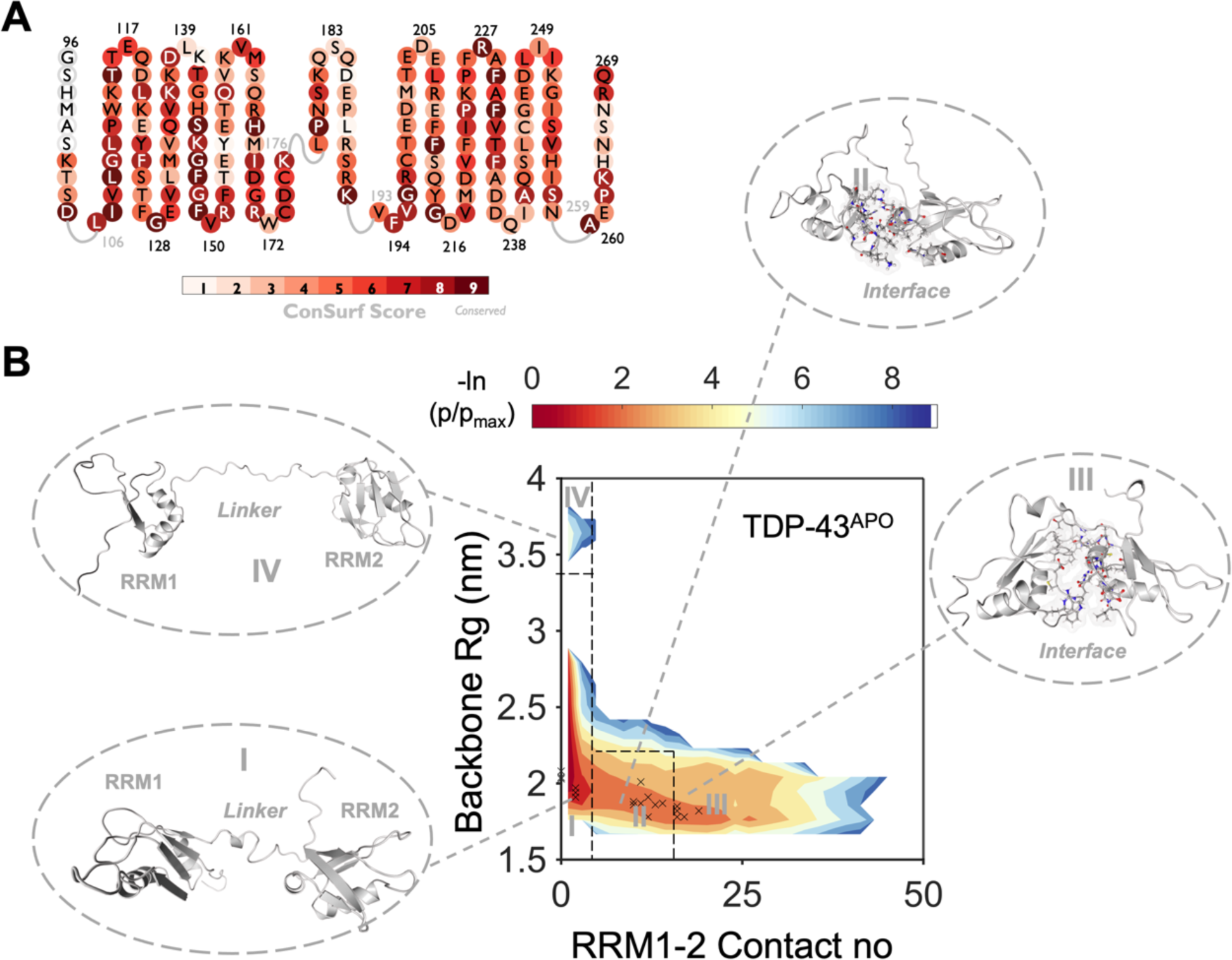
Characterization of the configurational space of RNA-free TDP-43. **A** Conservation score (See Methods) of TDP-43 amino acid sequence spanning K102-Q269. **B**. The two-dimensional free energy landscape of RNA-free TDP-43 is constructed using order parameters of contacts formed between RRM1 and RRM2 and radius of gyration of protein backbone atoms. Inter-RRM contacts and R_g_ values for the initial configurations that are used for production runs are marked by crosses. Representative structures for each basin are shown in ribbon representation.

### Tandem RRMs exhibit intrinsic conformational plasticity in the absence of RNA

Free energy landscapes (FELs) computed based on one or more structural descriptors can provide useful insights into the conformational landscape of flexible biomolecules [60]. We constructed FELs of tandem RRMs configurations obtained from the simulation trajectories to determine the structural changes which occur upon its interaction with RNA. In this pursuit, we chose two fundamental metrics: (i) the radius of gyration (R_g_), which gives insights into the spatial extent of protein backbone atoms; (ii) the number of contacts established between the RRM domains (**Table S2)**. By doing so, we aimed to describe the conformational landscape in terms of the interplay between compactness and the formation of the RRM1-RRM2 interface.

The conformer populations observed in the FEL were categorized into four distinct regions for the RNA-free ensemble. Representative structures of these populations are presented in **Figure 1B**, and the corresponding values of order parameters are given in **Table S3.** Within these regions, regions I and IV contain extended configurations that are devoid of RRM1-RRM2 contacts. Interestingly, region IV populates configurations which exhibit a partial unfolding of the RRM1 domain, resulting in a significant increase in the radius of gyration (R_g_) and loss of secondary structure profile (**Figure 1B, S3**). Upon binding with RNA, conformational fluctuations of the ensemble decreased (**Figure 2A**). Notably, the RNA-bound ensemble does not populate extended configurations found in basin IV. This observation indicates that RNA functions as a protective factor, preventing unfolding of TDP-43 by restricting the adoption of extended configurations devoid of RRM1-RRM2 interactions. Furthermore, region II in the RNA-free FEL emerges as the most densely populated microstate within the RNA-bound FEL. This basin corresponds to the RNA-bound NMR ensemble, characterized by an R_g_ value of 1.72 nm and 24 contacts between RRM domains. Considering that the initial configurations for both the RNA-free and RNA-bound simulations were the same, our findings emphasize the role of RNA binding in stabilizing the tandem arrangement of the RRM domains.

**Figure 2.**
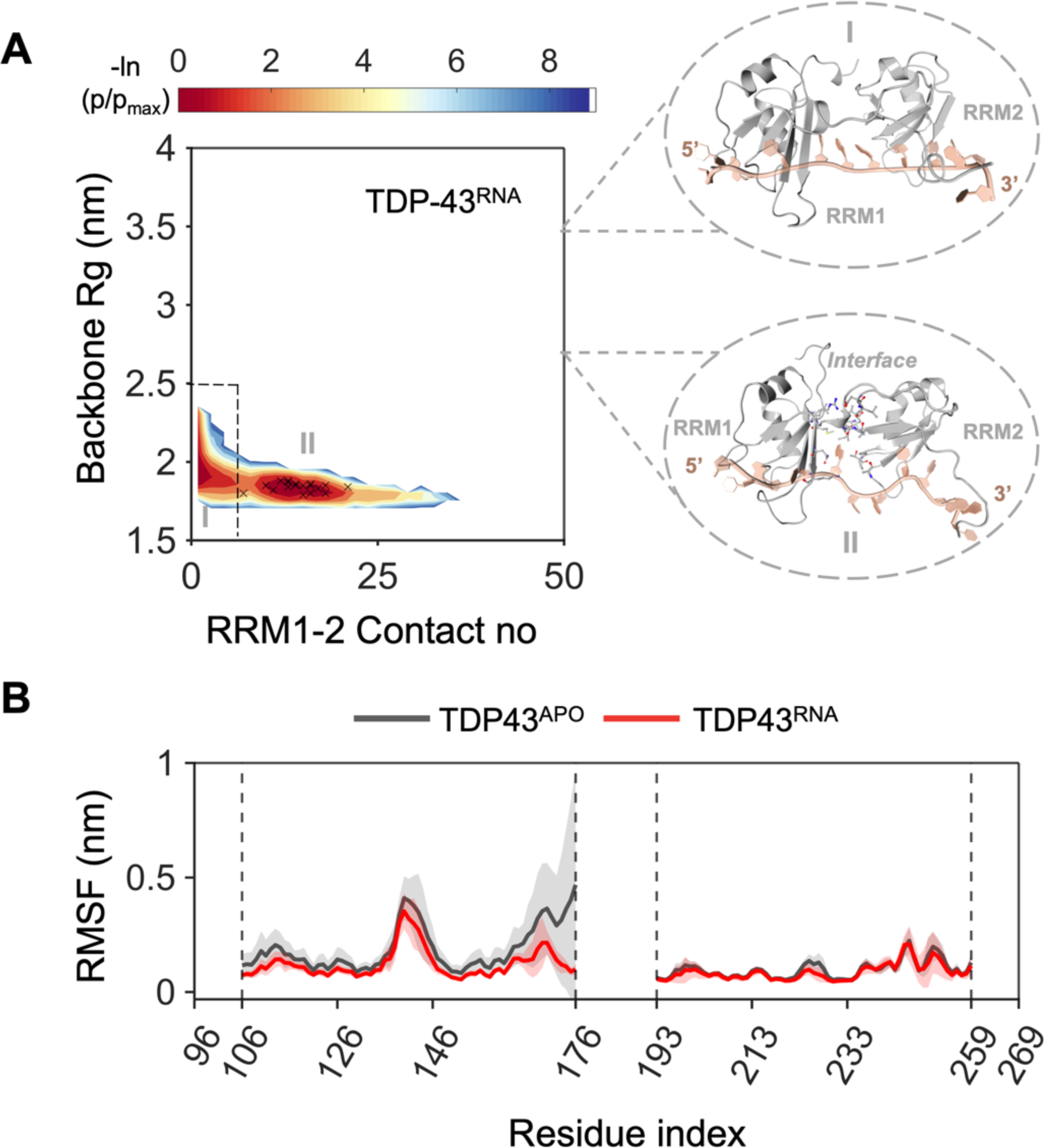
Characterization of the configurational space of TDP-43 RNA-bound state. **A.** Free energy landscape of RNA-bound TDP-43. RRM contacts and R_g_ values for the initial configurations that are used for production runs are marked on the surface. **B.** Per-residue RMSF values in RNA-free and RNA-bound states highlight the conformational flexibility of loop 3 and β4/5 region of the RRM1 domain in the RNA-free state.

Next, we pursued a more in-depth investigation to further explore the changes in the stability of the RRM domains upon RNA binding. To achieve this, we computed the root mean square deviations (RMSD) of the Ca atoms within the isolated RRM1 and RRM2 domains of the RNA-free and RNA-bound tandem constructs. Since RRM1 has a longer, flexible loop region (aa: K137-G146^RRM1-loop3^) compared to RRM2 (aa: I222-A228^RRM2-loop3^), we calculated RMSD by excluding all the loop regions from the RRM domains. Collectively, our results provide a clear depiction of the consistently higher stability found in the RRM2 domain when compared to RRM1 across the two states of TDP-43 (**Figure S4&S5**). RRM1 displays intrinsic conformational fluctuations in the absence of RNA, which stabilizes once RNA is bound. In contrast, RNA binding has no substantial impact on RRM2 stability. Therefore, tandem RRM domains have distinct stability profiles and their responses to RNA interactions.

The analysis of root mean square fluctuation (RMSF) for the Ca atoms within the RRM domains provides further insights into the individual contributions of each residue to towards conformational fluctuations, as depicted in **Figure 2B**. β-1, Loop-1, Loop-3, and β4/5 regions within RRM1 demonstrate increased flexibility. On the other hand, residues within RRM2 exhibit more rigid profile with minor fluctuations, primarily concentrated around the Loop-3 region (**Figure S6**). Overall, the stability analyses provided here elucidates the structural elements within the RRM domains that are prone to fluctuations.

### Inter-domain contacts play role in RNA binding while intra-domain contacts are preserved

Next, our aim was to gain a comprehensive understanding of how individual residues contribute to the alterations in the conformational landscape induced by the binding of RNA. For that purpose, we constructed protein-protein contact maps of the RNA-free/bound ensembles from MD simulations as well as NMR ensemble (**Figure 3A**).

**Figure 3.**
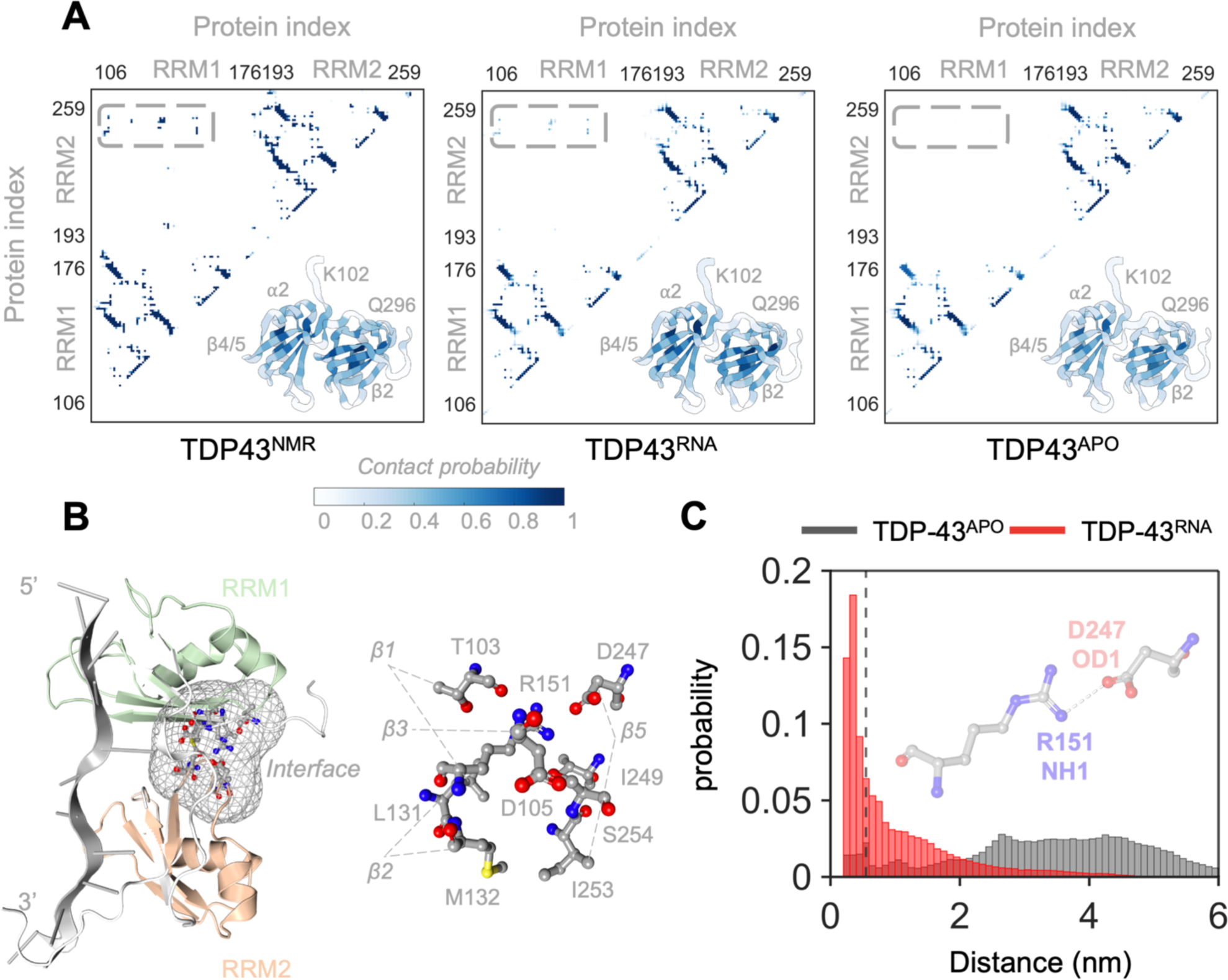
Intra-molecular interactions of TDP-43. **A**. Protein-protein contact maps of NMR, RNA-free, and RNA-bound ensembles. **B.** RRM1 and RRM2 contacts in RNA-bound ensemble shown on the representative structure of region II in RNA-bound free energy landscape. **C.** Distance distribution of R151-D247 representing the inter-RRM salt bridge formation. Bars within 0.6 nm as indicated by dashed vertical line correspond to salt-bridges.

First, we compared contact maps from the NMR ensemble with those from the RNA-bound ensemble. The most notable distinction was the weakening of the interactions between RRM1 and RRM2 over the simulation duration, suggesting a higher degree of plasticity within the RRM interface. Similarly, linker residues (aa:177-191) exhibit reduced contacts, consistent with the structurally disordered nature of this region. Moreover, we conducted fraction of native contacts analysis, which provided further evidence that the RNA-bound ensemble preserves intra-RRM domain contacts and exhibits dynamic inter RRM domain interactions (**Figure S7**). Collectively, these results indicate that the RNA-bound ensemble shows greater flexibility in the positioning of RRM domains relative to each other while maintaining the RNA in the binding pocket. The overall preservation of RRM1-RRM2 contacts indicates their significance in the context of RNA binding (**Figure 3B**).

Next, we compared the contact maps between the RNA-free and RNA-bound ensembles. It was observed that intra-RRM domain interactions remain largely unchanged and maintain their structural integrity regardless of the RNA binding status for the ensemble averages. However, it is important to note that the basin with a partially unfolded RRM1 population is low (**Table S3**), causing its contribution to disappear when considering the ensemble average. On the other hand, RNA binding induces a closer proximity between the RRM domains, resulting in the formation of an inter-RRM interface. In contrast, the RNA-free ensemble does not exhibit any notable and/or stable contacts between the RRM1 and RRM2 domains. Upon RNA binding, specific residues within RRM1 in the β1, β2, and β3 regions, establish significant protein-protein contacts (with a probability of forming greater than 0.45) with residues in the β4/5 region of RRM2. Particularly, interactions such as T103-D247, D105-D247, D105-S254, L131-I249, M132-I253, R151-D247, R151-I249, and R151-S254 contribute to the formation of the interface between RRM1 and RRM2 (**Figure 3B**). Lastly, residues in the β4/5 of RRM1, show an increase in the intra-domain contacts in the presence of RNA (**Figure S8**).

As previously demonstrated [61], the inter-domain salt bridge between R151 in RRM1 and D247 in RRM2 is pivotal for the stability of TDP-43, and the destabilization of this salt bridge leads to impaired GU-rich RNA binding. To assess how the dynamic RRM1-RRM2 interface influences this salt bridge, we measured the distance between the nitrogen atom of the R151 and oxygen atom of the D247 and computed its probability distribution over all trajectories (**Figure 3C**). As evident from the distance distributions, the RNA-bound state shows a stable R151-D247 salt-bridge. On the other hand, most of the configurations sampled in the RNA-free state have impaired R151-D247 salt-bridge. These results not only support the experimental findings but also highlight an inherent connection among the maintenance of the RRM1-RRM2 interface, the stability of the R151-D247 salt bridge, and RNA binding affinity (**Figure S9**).

### RNA dissociation from RRM domains does not depend on the tandem RRM arrangement

Previous studies on RNA binding to isolated RRM domains indicate that RRM1 exhibits higher binding affinity for GU-rich RNA compared to RRM2 [2, 5, 7]. Thermodynamic and nucleic acid binding experiments have led to the proposition that RRM2 may enhance RNA binding indirectly by stabilizing RRM1 [62]. Here, we investigated changes in the contact dynamics related to RNA binding upon the deletion of RRM1 or RRM2 to gain insights into the individual contributions of the RRM domains to RNA binding.

Deletion of RRM2 leads to a slight decrease in the overall stability of the RRM1 ensemble as indicated by the increased contribution of higher RMSD configurations towards the cumulative probability distribution (**Figure 4A**). On the other hand, upon the deletion of RRM1, partially unfolded RRM2 configurations with increased fluctuations become slightly more populated compared to the tandem RRM ensemble (**Figure 4A**). Residues within the β4/5 regions of the RRM1 and RRM2 display greater flexibility in these truncated ensembles (**Figure S10**). Since β4/5 region of the RRM2 is involved in interactions with the β1, β2, and β3 regions of RRM1 (**Figure 4B**), the increased flexibility can be attributed to the loss of these interactions in the truncated RRM2 ensemble. Additionally, the increased fluctuations of the β4/5 region of the RRM1 in the truncated RRM2 ensemble could be explained by its connectivity to the more flexible linker (**Figure S10**).

**Figure 4.**
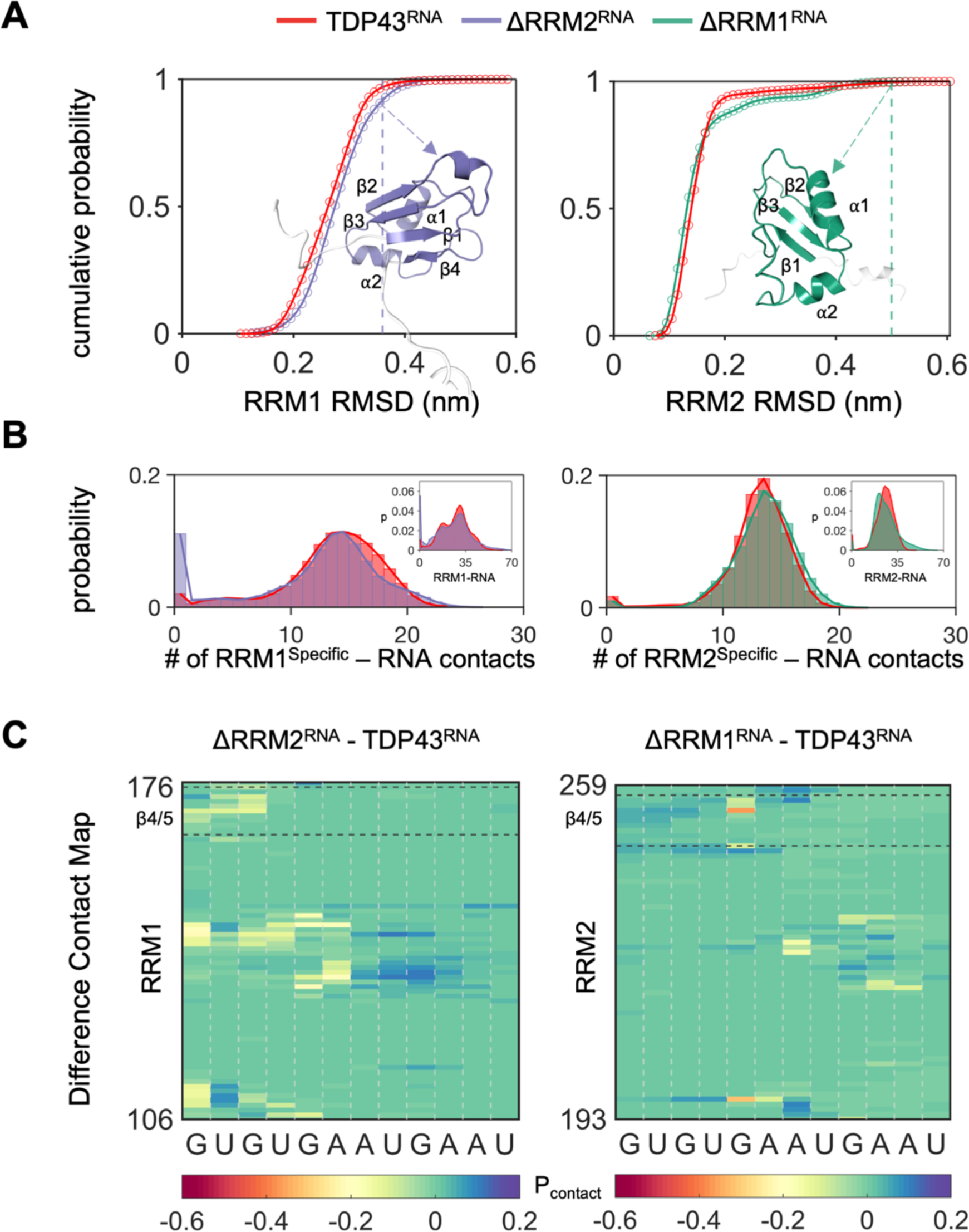
Individual domain contribution to the dissociation of RNA. **A.** Cumulative probability of RMSD for RRM1 and RRM2 domains with comparison to the constructs having either only RRM1 domain or RRM2 domain. Slightly increased RRM1 fluctuations upon the deletion of RRM2 and slightly enhanced probability of populating partially unfolded RRM2 upon the deletion of RRM1 are depicted. The last frames from the simulations started with the NMR representative (Model #0) are shown along with the RMSD values they have. **B**. Histogram of total RRM1-RNA contacts and RRM2-RNA contacts. Specific RRM-RNA contact probabilities are shown in the smaller panels, and **C.** Difference of the average probabilities for contacts between protein residues and RNA nucleotides. Positive values indicate increased contact probability and negative values indicate decreased contact probability in the constructs, ΔRRM1 and ΔRRM2.

Next, we investigated how RNA binding is affected by the loss of the tandem arrangement. For that purpose, we first evaluated the sum of contacts each domain formed with RNA (**Figure 5B**) and second, we constructed difference contact maps by subtracting the averaged contact probabilities of the tandem ensemble from the truncated ensembles (**Figure 4C**). Additionally, we plotted distributions of the specific contacts with RNA (**Figure 4B**).

**Figure 5.**
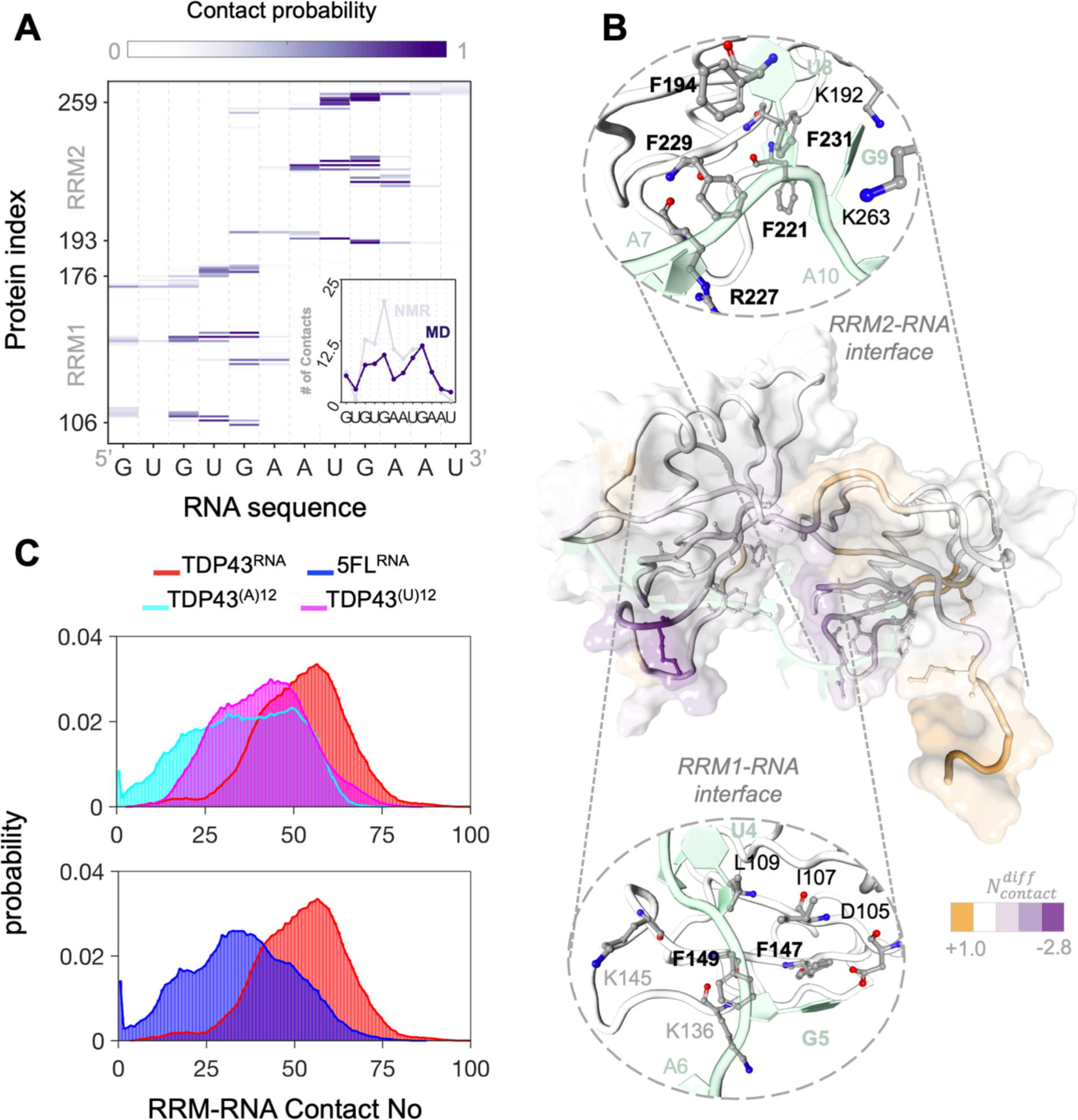
Dynamic interface between TDP-43 and GU-rich RNA. **A.** Protein-RNA contact map averaged over RNA-bound ensemble. Change in contact propensity of RNA nucleotides are shown in the small panel. **B.** Dynamic RNA binding mode observed in the MD simulations. The difference of contact propensity of the residue with RNA between MD ensemble and NMR ensemble is shown in the representative structure of region II in RNA-bound free energy landscape. RRM1-RNA and RRM2-RNA interfaces formed in the MD simulations are shown in the small panels. **C.** Distribution of the contact number between RRM domains and RNA for different constructs: proteins having GU-rich, polyA, polyU sequences, and 5FL mutant having GU rich RNA sequence.

According to our results, the loss of the RRM2 does not alter the overall distribution of the sum of contacts formed between RRM1 and RNA, and it does not affect the average contact propensity between RRM1 and RNA significantly (**Figure 4C**). However, there is a slight increase in the population of configurations with no contacts with RNA. Notably, the lost contacts are sequence-specific, particularly involving F147/F149 and G5 (**Figure 4B&C**). The loss of the RRM1 shifts the median of the distribution of the sum of the formed contacts between RRM2 and RNA. The decreased contact propensity, however, is not attributed to sequence-specific contacts, as interactions of conserved Phe residues with 3’ RNA remain intact (**Figure 4B&C**).

In summary, the average contact propensities between RRM domains and RNA remain largely unchanged, indicating that RNA dissociation is independent of the tandem RRM arrangement. However, the tandem RRM arrangement contributes to the conformational stability of each domain.

### GU-rich RNA interacts with RRM1 and RRM2 through highly conserved Lys and Phe residues

TDP-43 is known for its sequence-specificity towards GU-rich RNA. RRM1 has been demonstrated to have more affinity for GU-rich RNA than the RRM2 domain alone in the past [7]. Many independent studies [2, 5, 7] have collectively established that both domains are necessary to achieve high-affinity binding to the GU-rich motifs. The main interactions that retain GU-rich RNA in the binding pocket were proposed to be Phe147 and Phe149 [5]. Considering the prevailing notion of sequence specificity, it becomes important to assess the stability of interactions between protein residues and RNA nucleotides.

Initially, we conducted a comparative analysis of residue contributions to protein-RNA interactions observed in both NMR and RNA-bound MD ensembles (**Figure 4A, S11-13**). Both ensembles anchor RNA within the binding pocket through the stabilization of the 5’ end via interactions involving U4 and G5 with RRM1, and the 3’ end through interactions involving U8 and G9 with RRM2. Yet, we find a significant rearrangement in the protein-RNA contacts through MD simulations compared to the NMR ensemble. More precisely, RRM1-RNA contacts are less preserved in the MD ensemble compared to the NMR ensemble, while RRM2 residues exhibit stable interactions with RNA. Altogether, these observations indicate an intricate network of specific and non-specific (electrostatic) interactions involving the tandem RRMs and GU-rich RNA (**Figure 4A, S11-13**).

Next, we characterize the binding mode of the GU-rich RNA. Significant alterations were observed at the 5’ end of RNA and its interactions with RRM1. Specifically, interactions involving G3 and G5 experienced more fluctuations. Similarly, G5 departed away from S104, L106, K145, R151, D247, and V255, all of which were proximal in the reference NMR ensemble. In contrast, the 3’ end of RNA forms stable contacts with RRM2. Residues that contribute the most towards interactions with RNA are shown in **Figure 4B**. Many of these interactions are facilitated by conserved Phe and Lys residues (**Figure 1A**). F147-U4/G5, F149-G5, F194-U8, F221-G9, F229-U8/G9, and F231-U8/G9 interactions are well-preserved as indicated by an average contact probability greater than 0.85, depicting the crucial role of Phe residues in promoting binding specificity towards GU-rich RNA. Lysine-RNA interactions, on the other hand, are less preserved as many lysine residues are in disordered regions (**Figure S14**). Thus, it appears that stacking interactions play a significant role in preserving the sequence-specificity of RNA binding, while electrostatic interactions, primarily through RRM1, are of secondary importance.

Poly(U)_12_ and poly(A)_12_ sequences show reduced binding affinity towards TDP-43, with the latter exhibiting substantially weaker binding [2, 7]. Additionally, the 5FL mutant (F147/149/194/229/231/L) has been established as a control case for evaluating “impaired RNA binding” [63]. Therefore, to investigate the GU-rich specificity of TDP-43 further, we designed three additional simulation sets by changing the RNA sequence to poly(U)_12_ and poly(A)_12_, and by mutating the conserved Phe residues to Leu. We calculated the sum of the contacts between the RRM domains and RNA for each sampled configuration and computed their probability distributions (**Figure 4C**). The comparison of contact probability distributions indicates a significant loss of protein-RNA interactions for all the constructs which aligns with the experimental observations [2, 7, 63]. The shift is more pronounced for TDP43^(A)12^ and 5FL^RNA^ than the TDP43^(U)12^. Furthermore, this trend persists when considering only conserved Phe (F147, F149, F194, F221, F229, and F231) and RNA interactions for calculations (**Figure S15**) highlighting the importance of the stacking interactions with U4, G5, U8, and G9 for the sequence specific binding of the RNA to TDP-43.

In conclusion, our simulations elucidate the dynamic nature of RNA-RRM interactions which promote an overall stabilization of the tandem RRM arrangement. Nevertheless, we observed key differences between the binding modes of RRM1 and RRM2 towards GU-rich RNA which are linked to the relative enrichment of electrostatic versus hydrophobic interactions associated with either domain (**Figure S14**).

## DISCUSSION

Despite extensive research on the role of TDP-43 RRMs in pathological aggregation [3, 9, 13, 14, 25, 34, 35, 42, 44, 45, 62], achieving a comprehensive characterization of their structural dynamics in the presence and absence of nucleic acids based on available experimental approaches, has remained difficult. To address this gap, we employed an in-silico approach that utilized multi-replica, all-atom MD simulations, initiated from configurations taken from the NMR ensemble. This approach allowed us to explore the conformational heterogeneity of the RNA-free and RNA-bound states for functional TDP-43 (wild type, WT). Based on results provided here, we revealed that the RNA-free state exhibits a high degree of conformational plasticity, which is reduced upon RNA binding in the case of functional TDP-43. Altogether, our findings support the concept that ligand binding can alter the protein’s function by modifying the delicate equilibrium between states within the conformational landscape [64].

Consistent with previous observations, the binding of RNA enhances the stability of functional TDP-43 by stabilizing the β4/5 region within the RRM1 domain through intermolecular interactions with nucleic acid. In the absence of these interactions, the partially unfolded subbasin becomes accessible to TDP-43, potentially resulting in the accumulation of non-native TDP-43 in the cytoplasm. Thus, RNA binding not only plays a pivotal role in maintaining the functionality of the protein network but also acts as a protective factor, restricting the exploration of aggregation-prone microstates [65].

Previously, experimental and computational studies emphasized the high-stability of RRM2, and the possibility of an intermediate state in RRM2 that could trigger disease-related processes [41, 42, 62]. In our research, we observed that RRM2 remained highly stable and was unaffected by the absence of RNA. The studies proposing the existence of an intermediate transition state for RRM2 often characterized this state in the absence of RRM1. Our simulations, conducted without the presence of the RRM1 domain, also revealed one major stable peak and two relatively less stable minor peaks in the RRM2 RMSD distribution (**Figure S10**). These minor peaks may correspond to preceding configurations leading to the unfolding of RRM2. Overall, our findings suggest that RRM1 is stabilized by RNA, while the stability of the RRM2 is contingent on the presence of RRM1.

In addition to the intrinsic stability of RRM2 over RRM1 in tandem RRMs, GU-rich RNA establishes stronger contacts with RRM2 which are mediated by conserved Phe residues. While RRM1 has been shown to have higher affinity for GU-rich sequence than RRM2 domain alone, it can be argued that RRM1 plays a crucial role in facilitating the initial contact with the RNA through conserved Lys residues [7]. However, RRM2 is important in keeping the RNA within the binding pocket and maintaining the sequence-specific interactions, as it forms a greater number of contacts in this regard [66]. This observation also provides an explanation for why the tandem RRM domains collectively exhibit a higher affinity for RNA than either RRM1 or RRM2 alone.

In conclusion, all-atom MD simulations has enabled us to sample and explore various biophysical characteristics of RNA-free and RNA-bound states of TDP-43 RRMs. However, it is important to acknowledge several inherent limitations when simulating complex systems involving both a protein with disordered regions and unstructured RNA. These limitations include the accuracy of the RNA force fields and timescale constraints that prevent accessing all possible microstates [67, 68]. Additionally, all the initial structures used in simulations contained a functional RRM1-RRM2 arrangement due to the lack of RNA-free experimental structures. Despite these challenges, our findings illustrate that combining MD simulations with NMR experiments represents a powerful tool for investigating the dynamics of protein-RNA complexes [68, 69].

## SUPPORTING MATERIAL

Supporting material are given in Figure S1-15 and Table S1-5.

## AUTHOR CONTRIBUTIONS

B.O. conducted the simulations and performed the computational analysis. All authors designed the research and wrote the manuscript. J.M. supervised the research.

## ACKNOWLEDGEMENTS

This work was supported by NINDS and NIA R01NS116176. All-atom MD simulations were conducted with the advanced computing resources provided by Texas A&M High Performance Research Computing.

## DECLERATION OF INTERESTS

The authors declare no competing interests.

