## Supplementary Information for "RNA-binding tunes the conformational plasticity and intradomain stability of TDP-43 tandem RNA recognition motifs"

**Choice of RNA force field: Ensembles sampled by χOL3 and DESRES RNA force fields does not show structural differences.**

Two recent RNA force fields – (i) χOL3 [1-4] and (ii) DESRES [5], were tested in combination with the AMBER03ws protein force field to determine if the protein-RNA complexes were stable in simulations and thereby suitable for subsequent analysis. The ensembles of RNA bound TDP-43 with χOL3 and DESRES force fields are compared with respect to their stability, frequency of the native contacts of the protein-RNA interface, and total contact number between protein and RNA. RMSD values show no difference for the stability of the RRM domains (**Figure S2**). The minor fluctuations are observed in the loop regions which are expected and can be discarded. The changes of the protein and RNA interface are measured with native contact analysis by using two different cut-offs. The trend in preserving the native contacts at the interface are the same for both ensembles. Lastly, the average number of the total protein-RNA contacts, RRM1-RNA contacts, RRM2-RNA contacts are calculated individually for each frame and compared with the corresponding average NMR model value. Both simulation sets show the most crowded populations have %20-%30 of contact loss at the interface. As a result, for unstructured UG rich RNA, the χOL3 and DESRES force fields sample from the same configurational space. Therefore, we decided to use χOL3 as the RNA force field for the remaining simulation sets.

**GU-rich RNA does not leave its NMR binding pose in μs time scale.**

Native contacts can be defined as contacts in the minimized structures. The fraction of native contacts, then, can be determined by comparing the distances for each atom pair i and j with the reference distances using the equation introduced by Best et al (2013) [6]. According to this, fraction of the native contacts for atom pairs i which is selected from heavy atoms of the isolated RRM domains and j which is selected from the heavy atoms of the sequence specific nucleotides (G3, U4, G5, U8, and G9) are calculated (**Figure S7**). During 1 μs long simulations (20 μs in total), sampled conformations have a high tendency to preserve their RRM2-RNA interactions while the fraction of contacts between RRM1 and RNA show much greater flexibility. The most populated conformations keep the majority of their RRM1/2-RNA contacts indicating the simulation protocol followed for this study produces stable trajectories and could be used for further analysis. Furthermore, the ensemble preserves intra RRM domain contacts with very high frequency and fails to maintain most of its inter RRM domain interactions. This signifies the importance of the remaining contacts between the inter RRM domains.

**Table S1.** Details of simulation sets.

| Name | Initial Structure | RNA | Accumulation Time |
|---|---|---|---|
| TDP-43$^{RNA}$ | TDP-43 tandem RRMs: K102-Q269 (with 6aa tag, G96-S101) | GU-rich | 20*1μs |
| TDP-43$^{APO}$ | TDP-43 tandem RRMs: K102-Q269 (with 6aa tag, G96-S101) | No RNA | 20*1μs |
| TDP-43$^{(A)12}$ | TDP-43 tandem RRMs: K102-Q269 (with 6aa tag, G96-S101) | (A)12 | 20*1μs |
| TDP-43$^{(U)12}$ | TDP-43 tandem RRMs: K102-Q269 (with 6aa tag, G96-S101) | (U)12 | 20*1μs |
| ΔRRM1 $^{RNA}$ | L177-Q269 | GU-rich | 20*1μs |
| ΔRRM2 $^{RNA}$ | K102-R191 (with 6aa tag, G96-S101) | GU-rich | 20*1μs |

**Table S2.** Initial Rg and RRM1-RRM2 contact no for each trajectory used in sampling RNA-free state of TDP-43.

| Trajectory | Initial Rg (nm) | Initial RRM1-RRM2 contact no |
|---|---|---|
| 1 | 1.87 | 14 |
| 2 | 1.78 | 16 |
| 3 | 1.78 | 17 |
| 4 | 1.87 | 14 |
| 5 | 1.78 | 12 |
| 6 | 1.88 | 10 |
| 7 | 2.01 | 11 |
| 8 | 2.04 | 0 |
| 9 | 1.87 | 11 |
| 10 | 1.86 | 13 |
| 11 | 2.08 | 0 |
| 12 | 1.91 | 2 |
| 13 | 1.94 | 2 |
| 14 | 1.85 | 16 |
| 15 | 1.82 | 16 |
| 16 | 1.91 | 12 |
| 17 | 2.03 | 0 |
| 18 | 1.86 | 10 |
| 19 | 1.82 | 19 |
| 20 | 1.97 | 2 |

**Table S3.** Basins observed in the free energy landscape of TDP-43$^{Apo}$. The conformational ensemble that are representing each basin are defined using Rg and RRM1-RRM2 contact no.

| Basin | # of contacts | Rg (nm) | Representative | Size (total: 200000) |
|---|---|---|---|---|
| I | 0-5 | 2-2.8 | 2.37 nm, 0 contact | 113683 |
| II | 10-20 | 1.8-2.1 | 1.97 nm, 15 contacts | 13353 |
| III | >25 | 1.8-1.85 | 1.83 nm, 27 contacts | 807 |
| IV | 0-5 | >3.5 | 3.71 nm, 0 contact | 281 |

**Table S4.** Initial Rg and RRM1-RRM2 contact no for each trajectory used in sampling RNA-bound state of TDP-43.

| Trajectory | Initial Rg (nm) | Initial RRM1-RRM2 contact no |
|---|---|---|
| 1 | 1.85 | 14 |
| 2 | 1.88 | 13 |
| 3 | 1.80 | 18 |
| 4 | 1.85 | 10 |
| 5 | 1.79 | 15 |
| 6 | 1.85 | 16 |
| 7 | 1.88 | 12 |
| 8 | 1.84 | 21 |
| 9 | 1.85 | 15 |
| 10 | 1.84 | 21 |
| 11 | 1.87 | 13 |
| 12 | 1.82 | 11 |
| 13 | 1.86 | 16 |
| 14 | 1.85 | 18 |
| 15 | 1.84 | 13 |
| 16 | 1.83 | 17 |
| 17 | 1.80 | 16 |
| 18 | 1.86 | 14 |
| 19 | 1.80 | 7 |
| 20 | 1.85 | 14 |

**Table S5**. Basins observed in the free energy landscape of TDP-43$^{RNA}$. The conformational ensemble that are representing each basin are defined using Rg and RRM1-RRM2 contact number.

| Basin | # of contacts | Rg (nm) | Representative | Size (total: 200000) |
|---|---|---|---|---|
| I | 0-5 | 1.8-2.3 | 1.9 nm, 0 contact | 80420 |
| II | 7-22 | 1.75-1.95 | 1.87 nm, 13 contacts | 102505 |

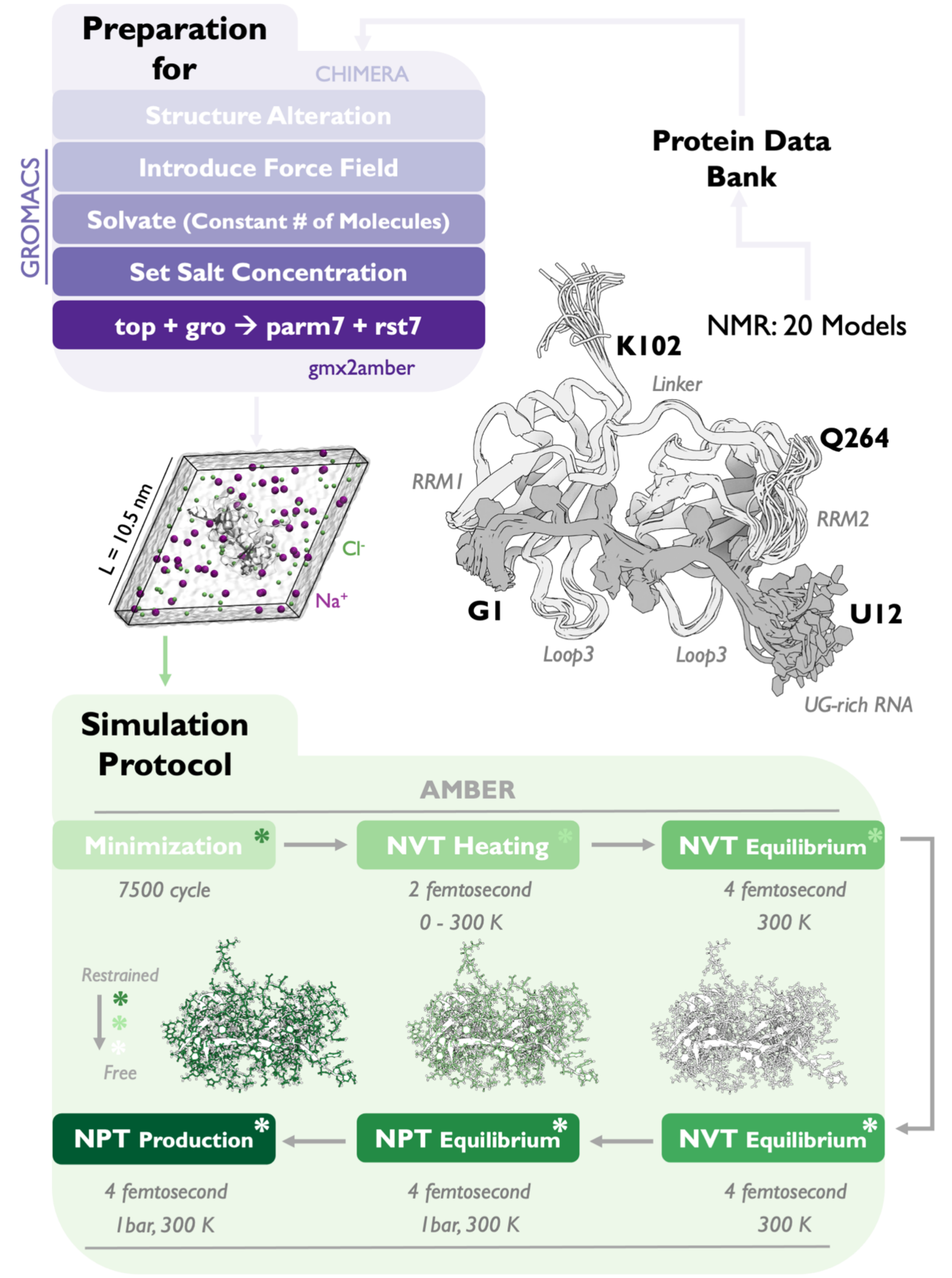

**Figure S1. Workflow for protein-RNA simulations**. NMR models of TDP-43 tandem RRMs with GU-rich (PDB ID: 4BS2) are used for demonstration.

A

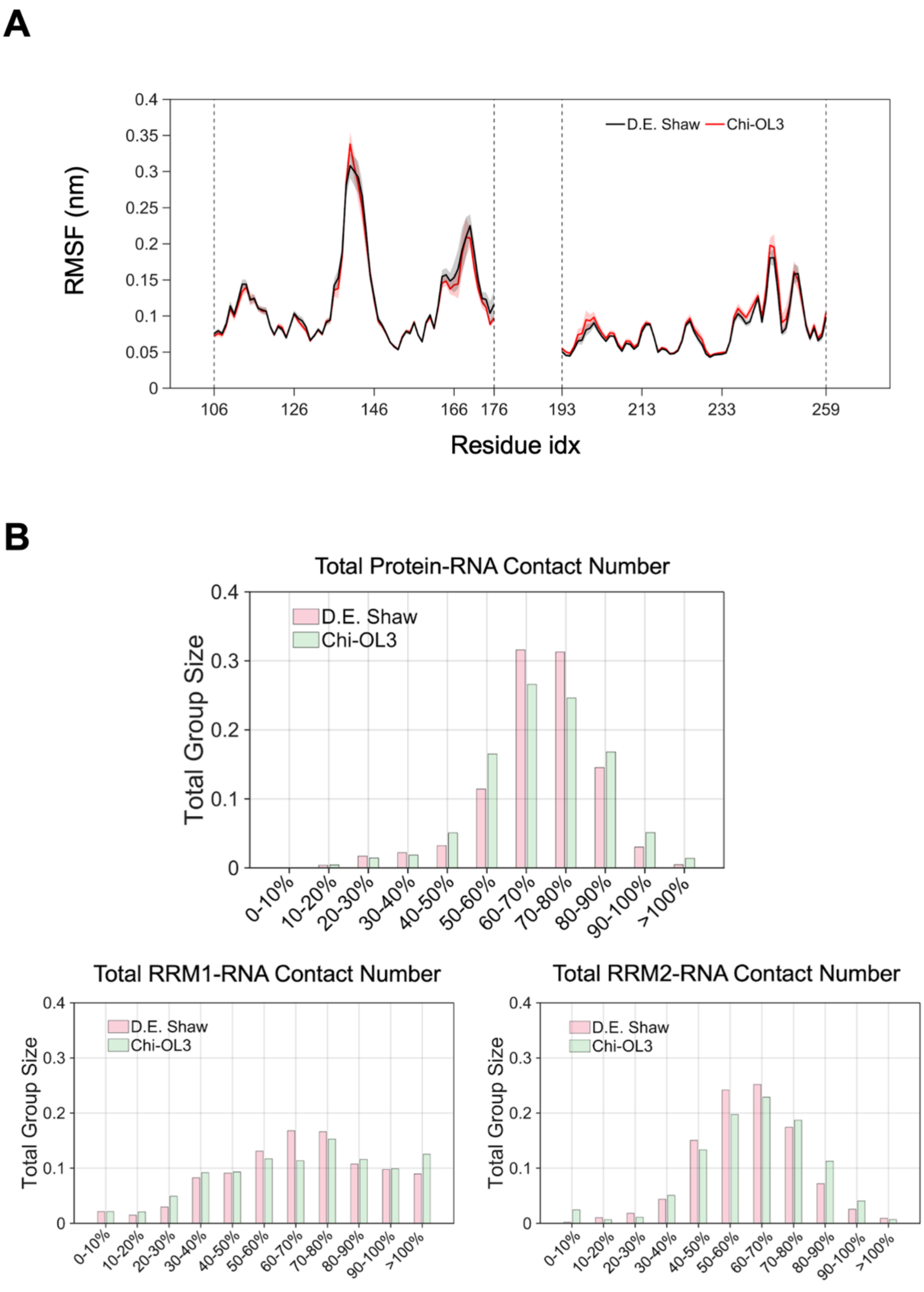

**Figure S2. χOL3 and DESRES force fields sample from the same configurational space.** **A.** RMSF values of individual RRM domains for ensembles sampled using different RNA force fields. **B.** Distribution of sum of formed contacts between protein and RNA. The value calculated for each configuration is compared with the average NMR contact number and then assigned to a group based on the percentage of preserving the contact number observed in NMR ensemble.

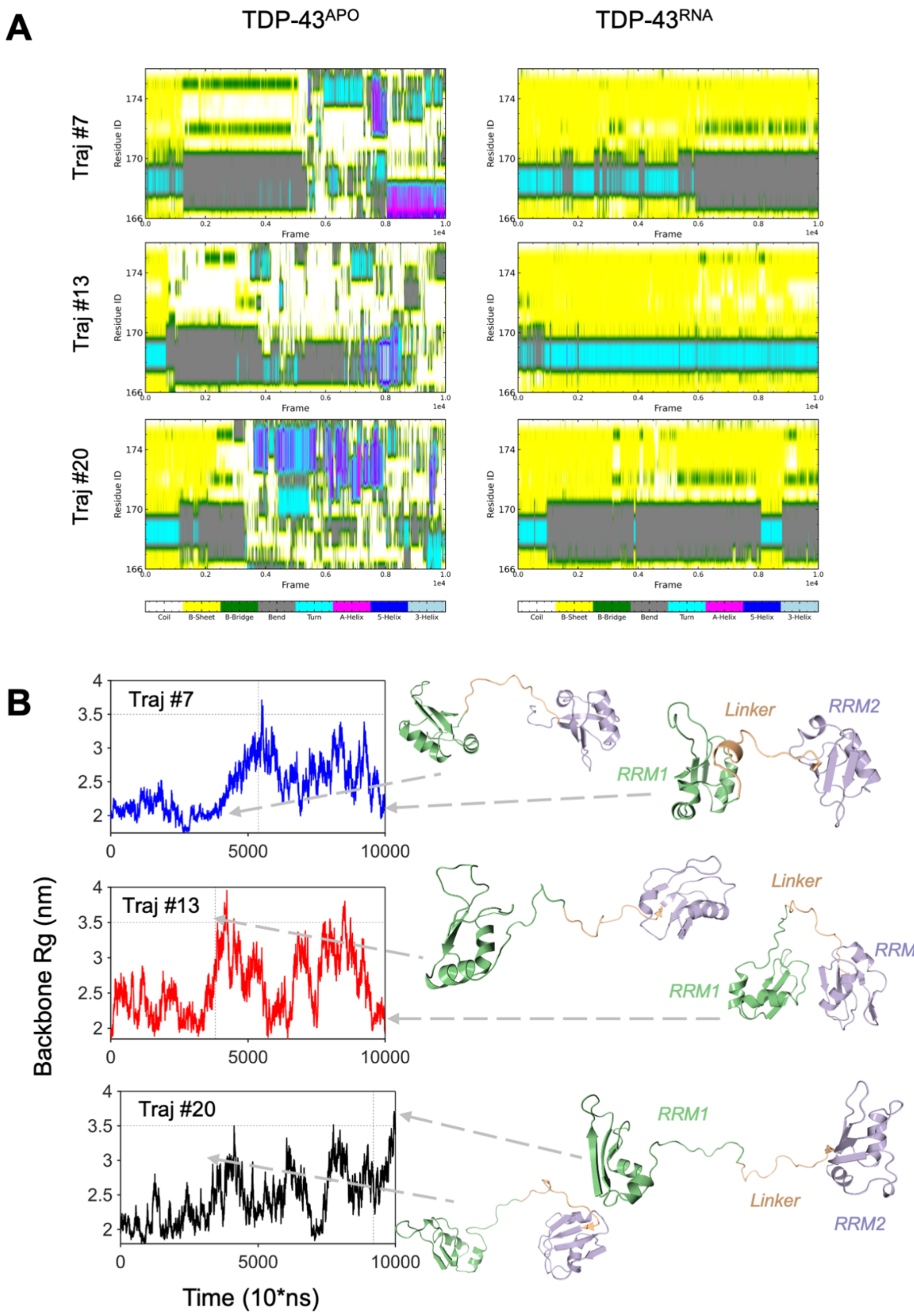

**Figure S3. RRM1 unfolding in the RNA-free TDP-43 ensemble. A.** Timewise secondary structure evolution of the β4/5 region of the RRM1 for the trajectories that constitute the basin IV of the RNA-free ensemble. **B.** Rg values of backbone atoms of the protein for trajectories that constitute the basin IV of the RNA-free ensemble.

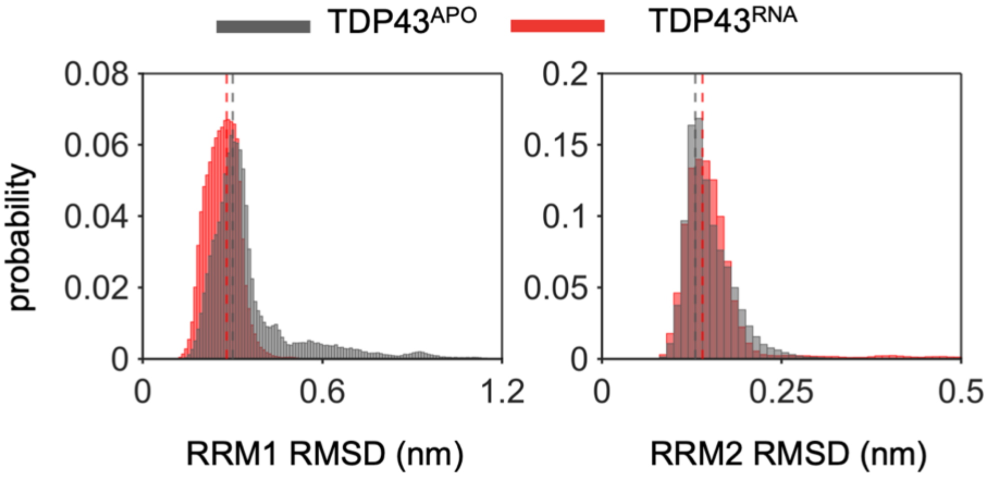

**Figure S4. Stability of individual RRM domains in the presence and absence of RNA.** Probability distributions of RMSD values calculated using Ca atoms of RRM1 and RRM2. Median values are shown as dashed lines.

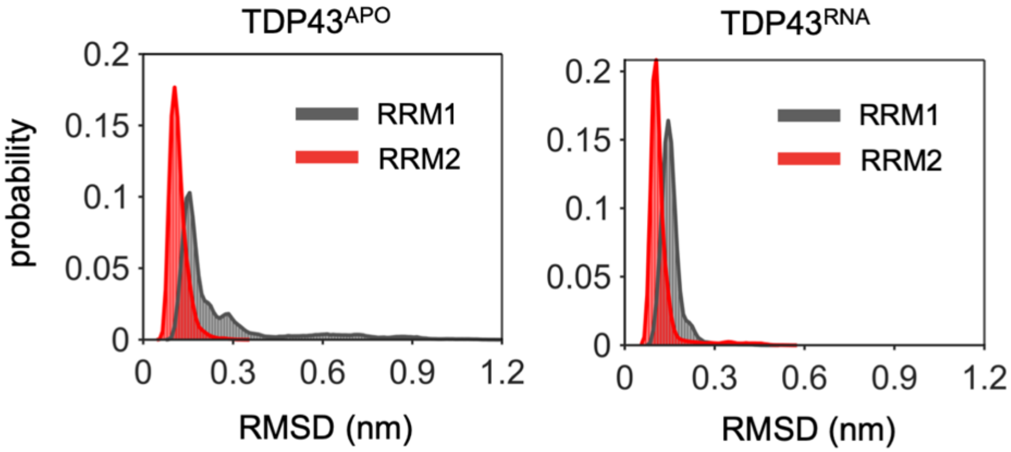

**Figure S5. Stability of folded regions in the presence and absence of RNA**. Probability distributions of RMSD values of Ca atoms which are reside in the folded regions of RRM1 and RRM2 domains. Folded regions are characterized by the visual inspection of the NMR structure, PDB 4BS2, having lowest energy.

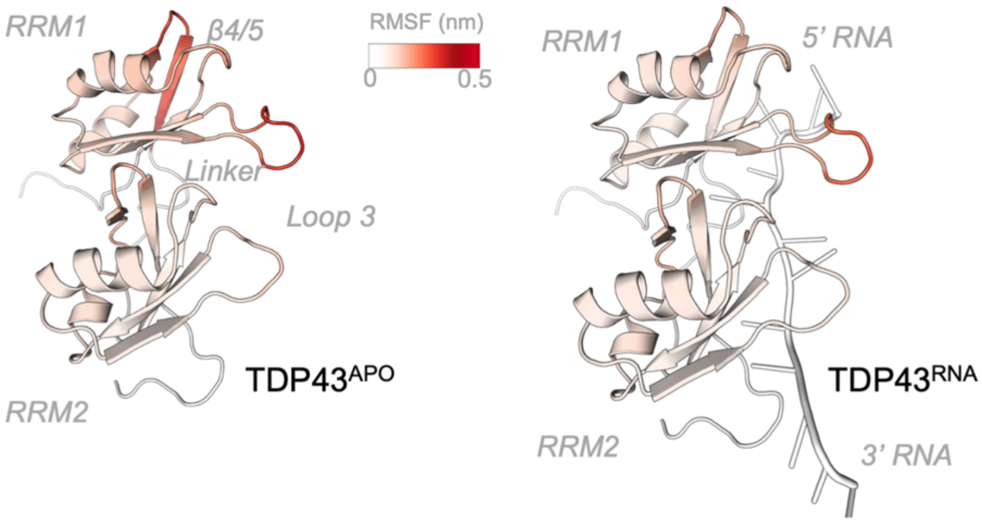

**Figure S6. Residue flexibilities of RNA-free and RNA-bound TDP-43**. RMSF values mapped on NMR structure (PDB: 4BS2) having lowest energy.

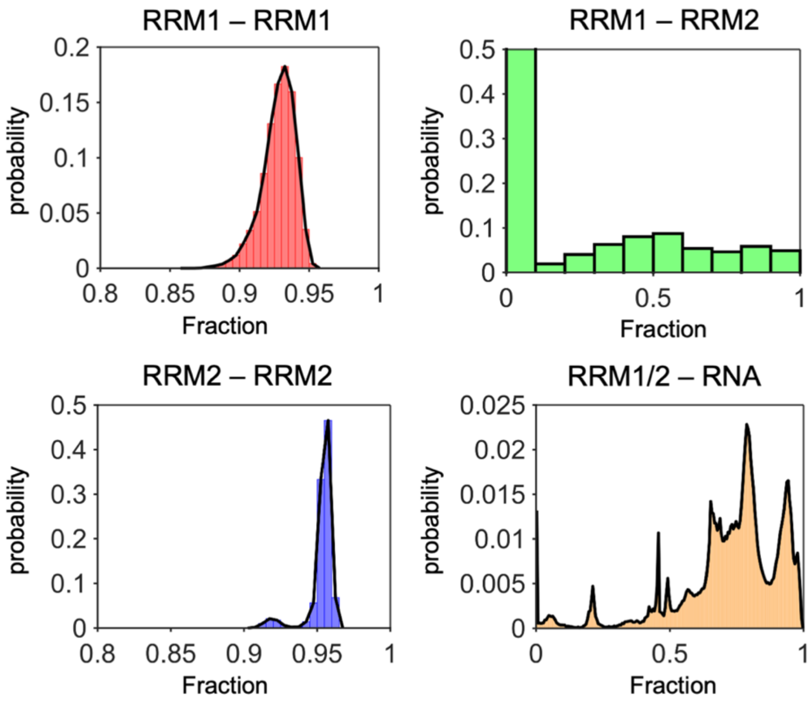

$$Q(r, r_o) = \frac{1}{1 + \exp(\beta(r - \lambda r_o)}$$

**Figure S7. Native contact analysis using soft cutoff method**. Only folded regions in the reference NMR structures are used for the calculations of fraction of native contacts. β is the softness of the switching function and is used as 5Å as proposed; while λ is the constant of reference distance tolerence with value of 1.8 for all atom simulations.

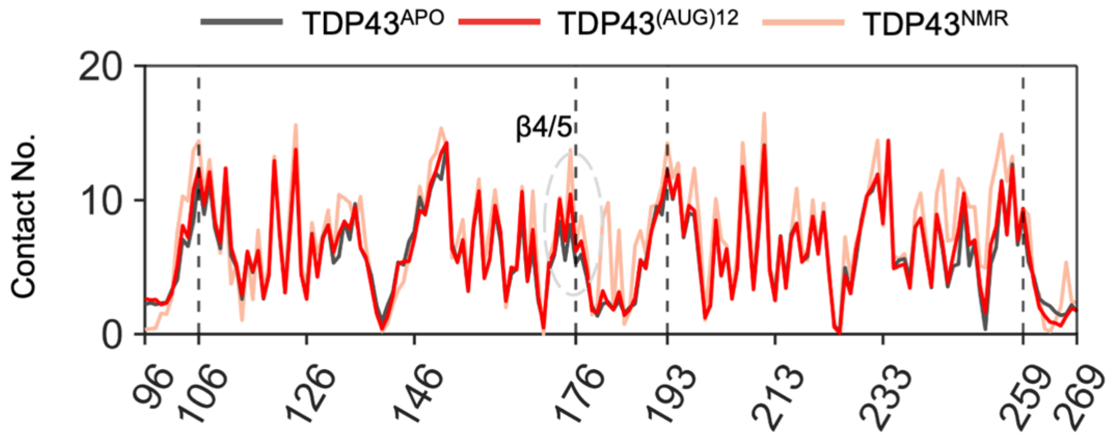

**Figure S8. Contact propensity of a residue with other residues in protein**. Values are averaged over 20 trajectory, and normalized over the frame number.

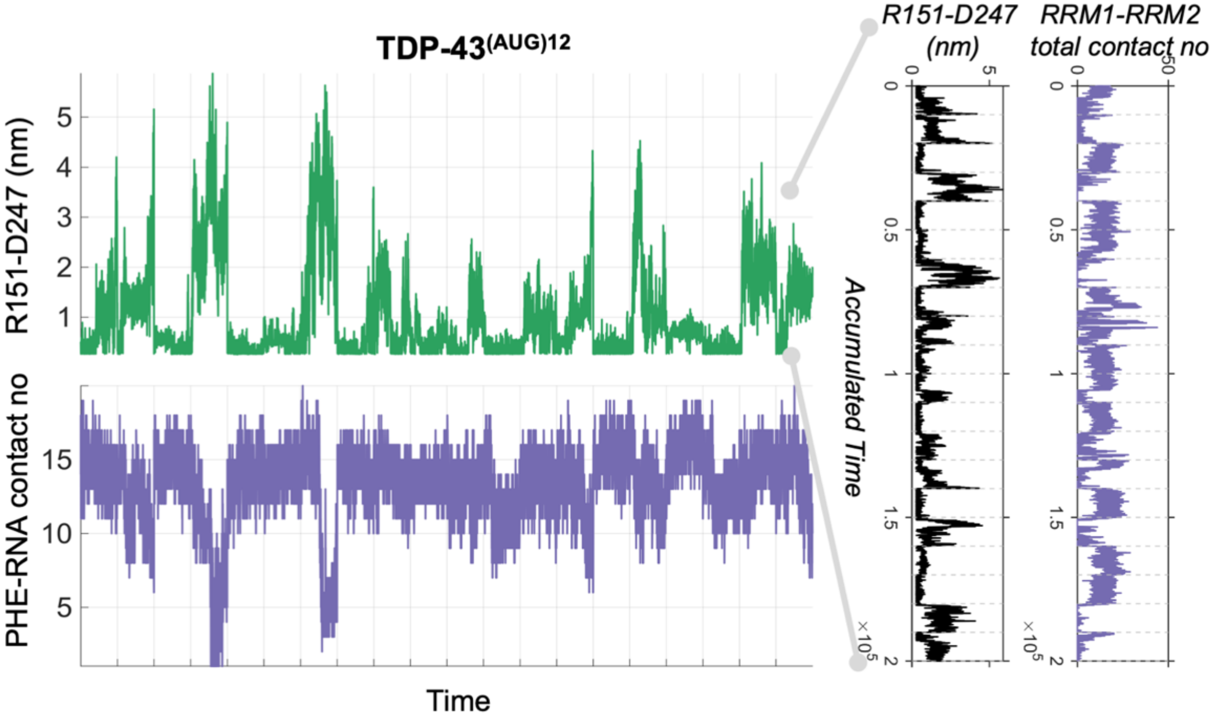

**Figure S9**. **Causality between PHE-RNA contact number, RRM1-RRM2 contact number, and R151-D247 distance.** The timewise distance of R151-D247 and sum of formed contacts between RRM1 and RRM2 and PHE residues and RNA are plotted for each 20 trajectory.

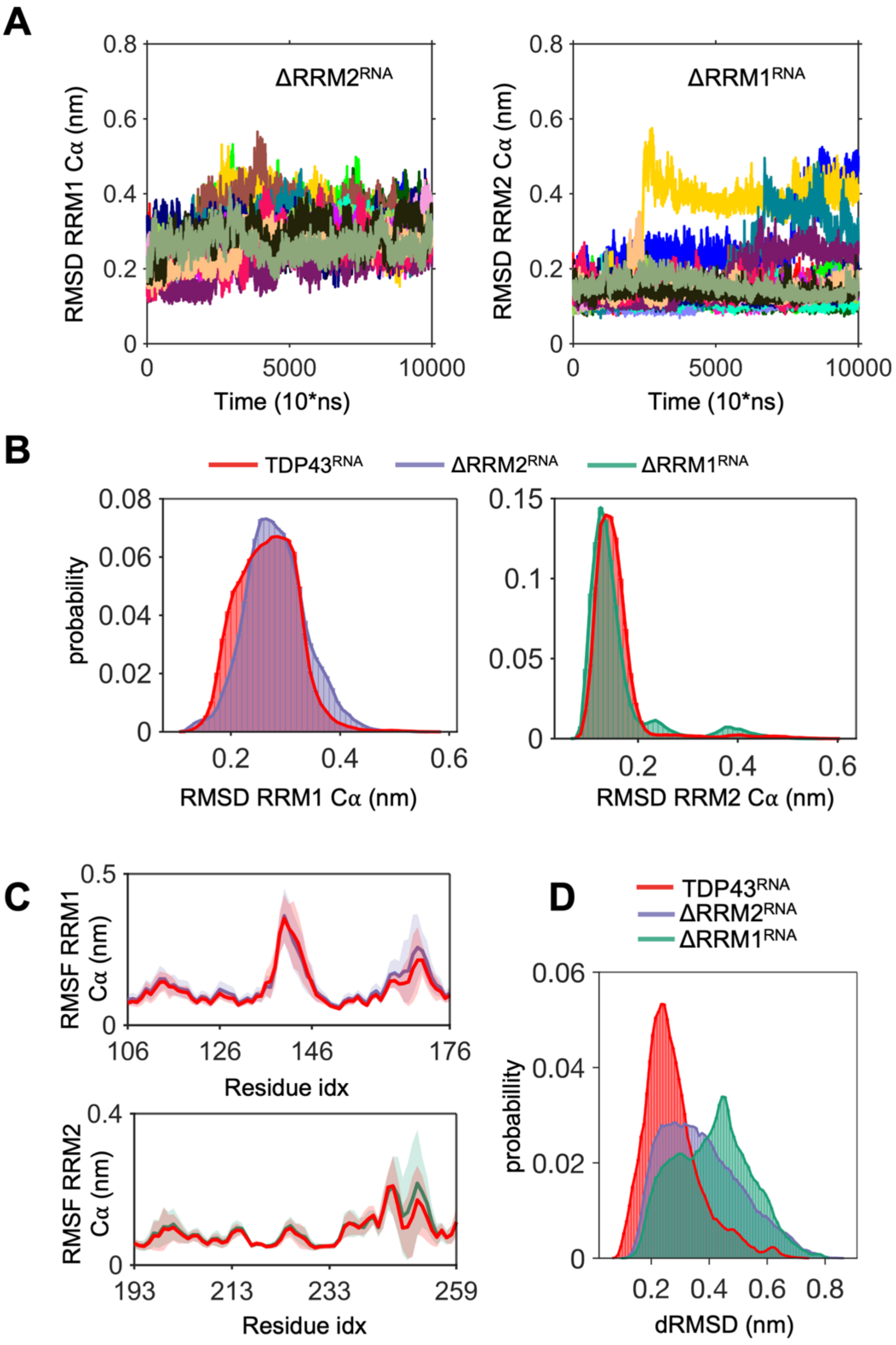
A
ΔRRM2^RNA
ΔRRM1^RNA
RMSD RRM1 Cα (nm)
RMSD RRM2 Cα (nm)
Time (10*ns)
0.8
0.6
0.4
0.2
0
5000
10000
B
TDP43^RNA
ΔRRM2^RNA
ΔRRM1^RNA
probability
0.08
0.06
0.04
0.02
0.15
0.1
0.05
RMSD RRM1 Cα (nm)
RMSD RRM2 Cα (nm)
C
RMSF RRM1 Cα (nm)
0.5
106
126
146
176
Residue idx
RMSF RRM2 Cα (nm)
193
213
233
259
Residue idx
D
TDP43^RNA
ΔRRM2^RNA
ΔRRM1^RNA
dRMSD (nm)

**Figure S10**. **Structural analysis of isolated RRM1 and RRM2 ensembles. A.** Timewise RMSD values for truncated RRM1 and RRM2., **B.** RMSD distribution of truncated RRM1 and RRM2 ensembles compared to the tandem RRM ensemble. RMSD analysis conducted using the Ca atoms within isolated RRM1 and RRM2 domains., **C.** Per-residue RMSF values in truncated ensembles., and **D.** Distribution of the distance of RMSD values of the Cα atoms reside in the linker region (L177-R191).

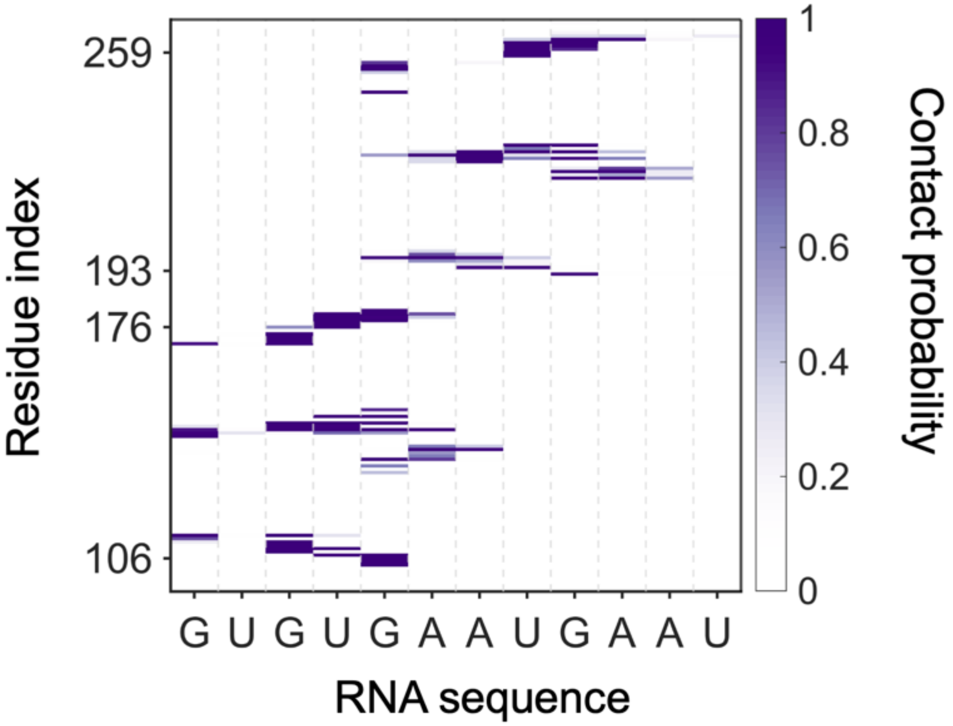

**Figure S11**. **Averaged normalized contact map of protein-RNA for NMR ensemble**. Contact maps are constructed for all 20 models, and contact propensities are averaged over 20 maps.

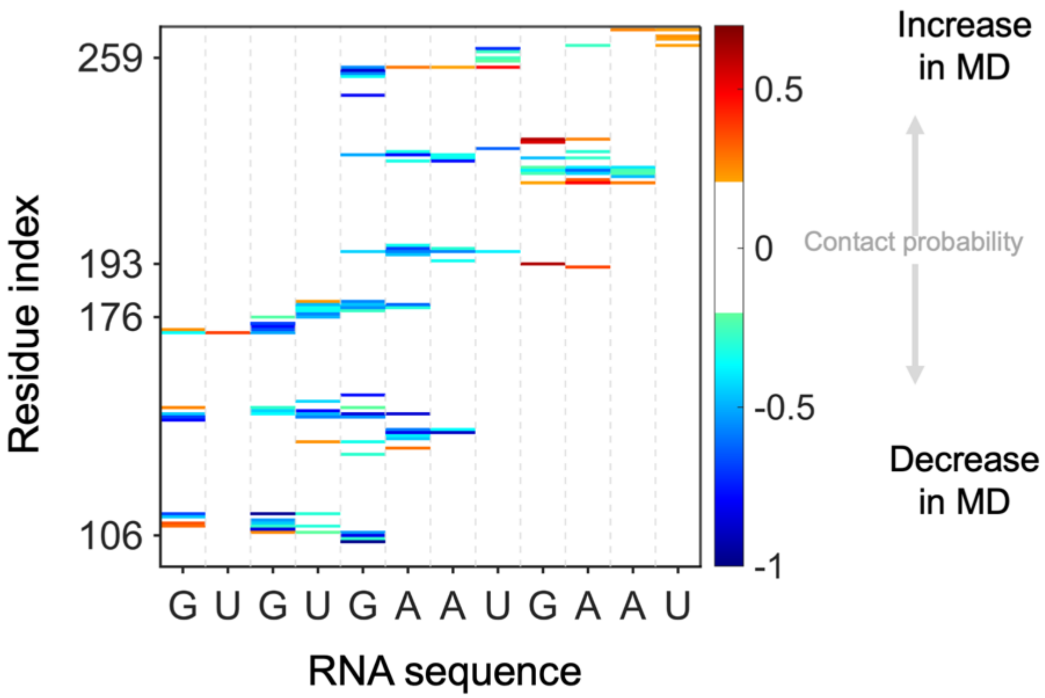

**Figure S12**. **Difference contact map for MD and NMR ensemble**. Normalized contact maps are constructed for all 20 ttajectories and then averaged over to obtain averaged contact map of MD ensemble. The contact probabilities of NMR ensemble, then, subtracted from the contact map of MD ensemble.

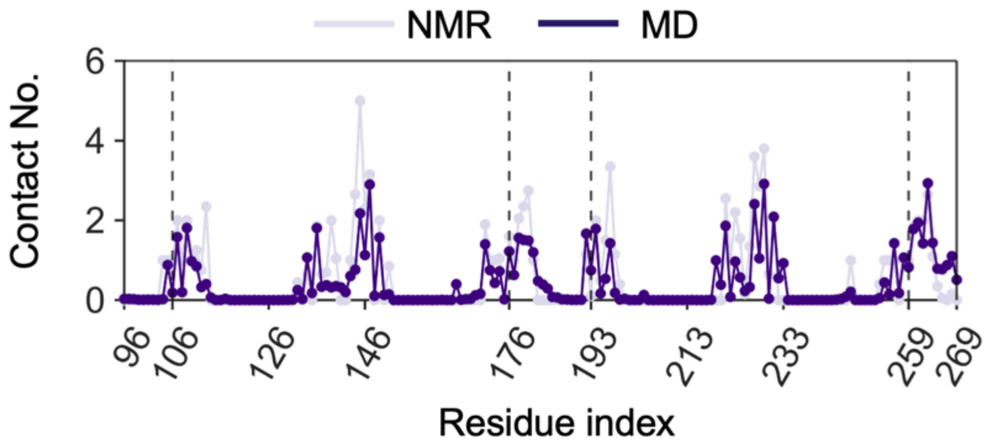

**Figure S13**. **Contact propensity of residues in MD and NMR ensembles.** Residuewise contact probabilities are obtained by summing the all possible contact probabilities of a residue. The plotted values are obtained by averaging the values of 20 trajectories/models.

**A**

**B**

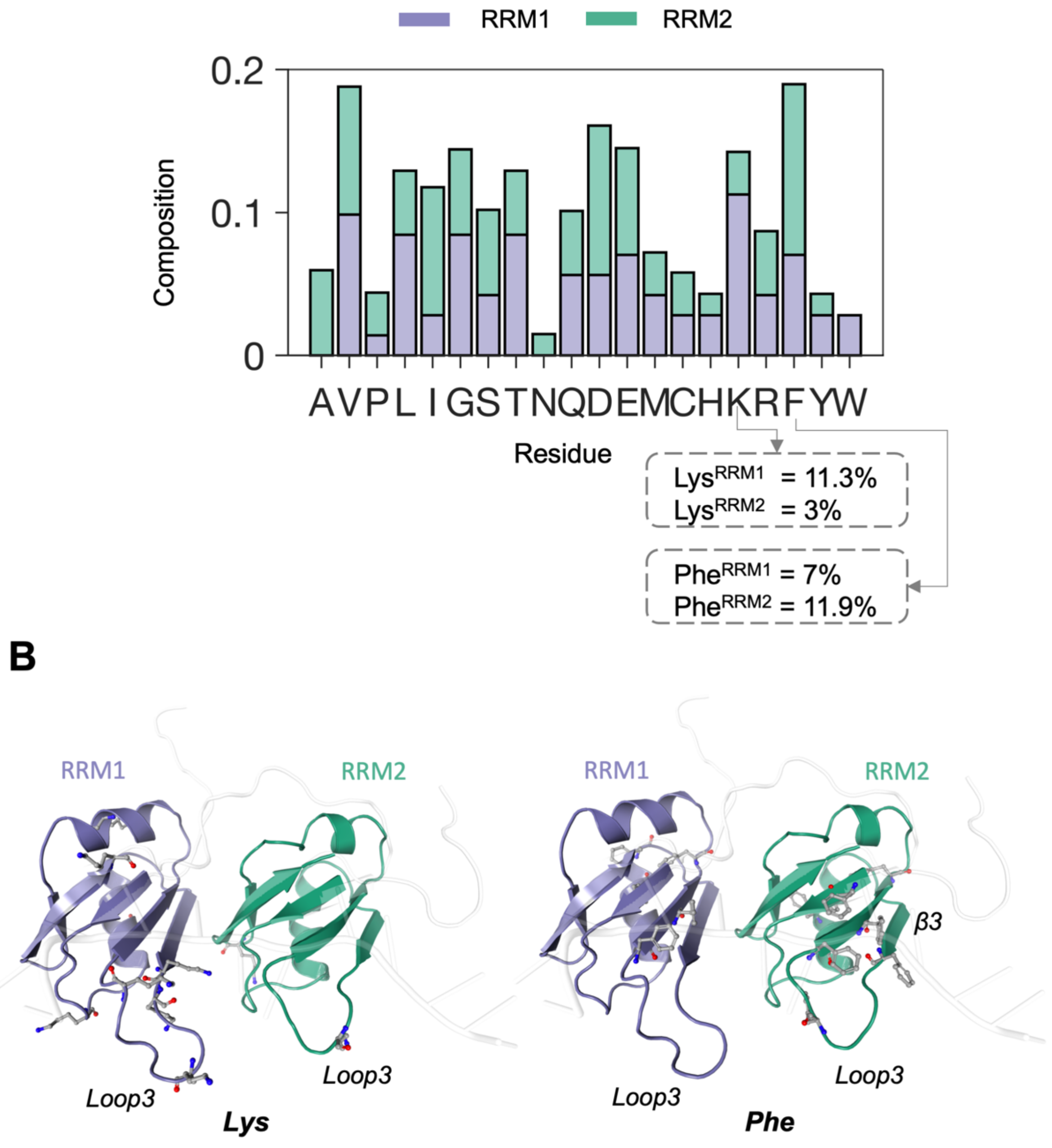

**Figure S14**. **Residue composition of the RRM domains**. A. The composition percentages of the Lys and Phe residues depict that RRM1 is richer in Lys residues compared to RRM2 while RRM2 is richer in Phe residues compared to RRM1. **B.** Lys and Phe residues are shown in the tandem RRM structure. Lys residues are clustered around loop3 of RRM1. Phe residues are clustered around β3.

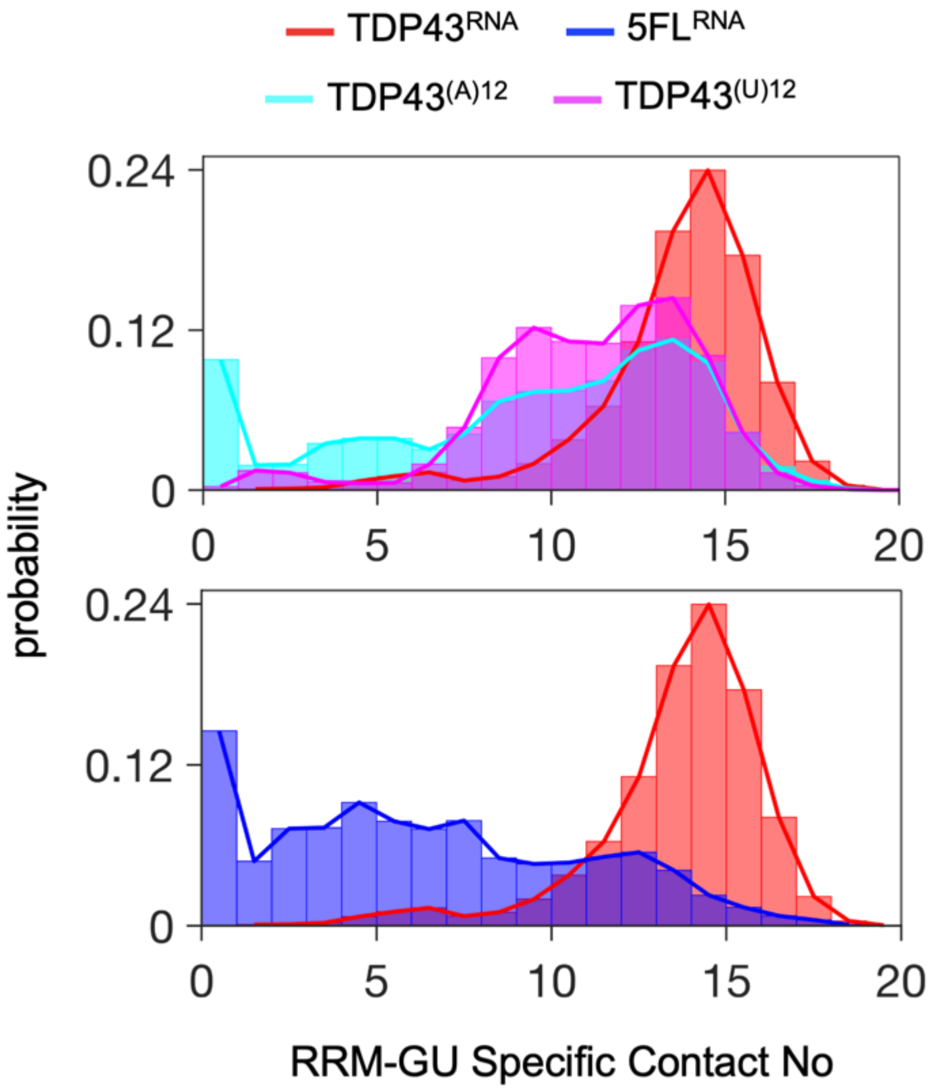

**Figure S15**. **Contact propensities of conserved PHE residues**. Histogram analysis of contacts formed between F147, F149, F194, F221, F229, and F231 residues or L147, L149, L194, L221, L229, and L231 residues in 5FL mutant and RNA having GU rich, polyA, or polyU sequences.
